# The Rhythm of Normality: A Comprehensive Normative Database for TMS-EEG Metrics with Reliability Characterization

**DOI:** 10.64898/2026.04.23.720403

**Authors:** Bruno Nascimento Couto, Enrico De Martino, Daniel Skak Mazhari-Jensen, Anne Jakobsen, Margit Midtgaard Bach, Alessandro Gianotta, Stian Ingemann-Molden, Thomas Graven-Nielsen, Adenauer Casali, Daniel Ciampi de Andrade

**Affiliations:** Center for Neuroplasticity and Pain (CNAP), Department of Health Science and Technology, Faculty of Medicine, Aalborg University, Aalborg, Denmark; Neural Engineering and Neurophysiology, Department of Health Science and Technology, Faculty of Medicine, Aalborg University, Aalborg, Denmark; Siena Brain Investigation and Neuromodulation Lab (Si-BIN Lab), Department of Medicine, Surgery and Neuroscience, Neurology and Clinical Neurophysiology Section, University of Siena, Italy; Institute of Science and Technology, Federal University of São Paulo, São Paulo, Brazil

**Keywords:** TMS-EEG, Normative Database, Test-retest Reliability, Cortical Excitability, Biomedical Signal Processing, Open Science

## Abstract

**Background:** Transcranial Magnetic Stimulation combined with electroencephalography (TMS-EEG) offers unique insights into cortical excitability and connectivity, yet current analyses are primarily limited to group-level inferences with little validation of individual reliability and feature redundancy.

**Objective:** To construct a comprehensive, open-access, and reliable normative dataset of TMS-EEG features that enables individual-level comparison

**Methods:** We aggregated TMS-EEG data recorded over the primary motor cortex (M1) from 164 healthy adults (30.8 ± 9.8 years; 88 female) across nine studies using harmonized acquisition and preprocessing pipelines. Reliability analysis was conducted on a test-retest subset (N=57) for 968 extracted features, evaluating systematic bias, absolute error, and relative reliability (Intraclass Correlation Coefficient categorized by the lower bound of the 95% confidence interval). Additionally, feature clustering was performed to quantify redundancy and correlations across the high-dimensional feature space. We then established normative distributions and developed an online benchmarking platform.

**Results:** Reliability analyses (N=57) of the high-dimensional feature set revealed that 525 out of 968 features (54.3%) met at least moderate reliability standards (ICC lower bound > 0.5). Cluster analysis indicated substantial redundancy among metrics, with three distinct clusters having a moderate-to-high internal correlation (|*r*| = 0.64, 0.48, 0.39, respectively). Finally, normative data from the database identified abnormal results in a test patient, supporting the feasibility of individual-level classification in an open-science framework.

**Conclusions:** Harmonization of data acquisition and analysis pipelines led to the development of a reliable normative M1 TMS-EEG reference. This publicly available resource provides a validated tool for future individual-level classifications and an open platform for ongoing community contributions.

## 1. Introduction

The combination of transcranial magnetic stimulation (TMS) and electroencephalography (EEG) provides a powerful approach for non-invasive interrogation of human cortical physiology (Ziemann et al., 2026). In this paradigm, a brief magnetic pulse generated by the TMS coil induces localized electrical currents in the cortex, while EEG captures the ensuing neural responses with millisecond precision (Chung et al., 2015; Ilmoniemi et al., 1997). Grounded in this “perturb-and-measure” framework, the TMS-EEG method allows for the quantification of cortical excitability, mapping of effective connectivity through the propagation of TMS-evoked potentials (TEPs), and characterization of large-scale network dynamics (Momi et al., 2023). Evidence indicates that TMS-EEG can detect altered excitability and connectivity patterns in groups of patients across a range of neuropsychiatric and neurological conditions, including stroke (Sarasso et al., 2020, 2025), Parkinson’s disease (Pei et al., 2022), disorders of consciousness (Casarotto et al., 2016), pain (De Martino et al., 2023, 2024), major depressive disorder (Strafella et al., 2022), and schizophrenia (Ferrarelli & Phillips, 2021).

The clinical translation of TMS-EEG remains limited by two principal challenges. The first is methodological, where heterogeneity in TMS hardware, stimulation parameters, artifact suppression, and EEG preprocessing pipelines are obstacles to reproducibility and comparability across studies (Belardinelli et al., 2019; Brancaccio et al., 2024). The second is neurophysiological: the TMS-evoked response is modulated by the instantaneous state of the brain, including wakefulness state and ongoing oscillatory activity, which introduces substantial variability (Guzmán López et al., 2022; Momi et al., 2023). This state dependence is analogous to other physiological measures, such as heart rate or blood pressure, which fluctuate over time and across individuals, but yet retain diagnostic value when deviations exceed normative thresholds (Sandercock, 2007). It remains unclear if the magnitude of TMS-EEG feature variability reflects intrinsic neural variabilities, and if it is proportional to genuine pathophysiological alterations associated with brain disorders (Cao et al., 2021; Tremblay et al., 2019). Additional sources of variability, including participant-related factors such as age and sex, as well as sensory confounds from auditory and somatosensory inputs, further complicate the interpretation of genuine cortical responses (Conde et al., 2019; Farzan & Bortoletto, 2022; Fecchio et al., 2025).

Clinical translation of any biomarker requires demonstrated reliability, or the capacity to yield stable measurements under unchanging conditions, which is a prerequisite for validity. This includes relative reliability for diagnostic use and absolute reliability for monitoring change over time (Aronson & Ferner, 2017; Hartmann et al., 2023). In this context, TMS-EEG features must demonstrate sufficient temporal stability and intersession reliability to support their use as biomarkers of cortical function (Bertazzoli et al., 2025; Kerwin et al., 2018).

While several studies have assessed TEP reliability (Belardinelli et al., 2019; Bertazzoli et al., 2021; Casarotto et al., 2010; Casula et al., 2021; De Goede et al., 2020; Guidali et al., 2023; Kerwin et al., 2018; Lioumis et al., 2009; Mancuso et al., 2021; Ozdemir et al., 2021a, 2021b; She et al., 2024; Ter Braack et al., 2019), the literature lacks a systematic, large-scale approach including a large number of TMS-EEG excitability and connectivity metrics classification (Bertazzoli et al., 2025), along with reference ranges that allows for individual participant interpretation. Normative databases provide this essential context: the range of healthy variation, against which an individual’s measurement can be compared to determine if it deviates from the norm (Marquand et al., 2016; Thatcher & Lubar, 2023). The creation of an open normative database would provide the necessary framework for translating TMS-EEG features into individual-level clinical practice.

This study directly addresses these two challenges. First, we computed a comprehensive set of M1 TMS-EEG features from studies with similar data collection and preprocessing pipelines, and we conducted the largest systematic reliability analysis to date for these features. Finally, we establish a robust and openly available normative database of TMS-EEG features, demonstrating its utility by enabling individual-level comparison against healthy norms. This work represents a first attempt to move TMS-EEG from group-level research towards precision level application.

## 2. Methods

### 2.1. Participants

Data were aggregated from baseline assessments from nine previous studies centered on symptom-free healthy individuals. Eight studies were conducted at the Center for Neuroplasticity and Pain (Aalborg University, Denmark) and one at Siena University (Italy). Assessment of test-retest reliability was based on a subset of participants who underwent two identical sessions separated by at least 72 hours. Key methodological aspects of each study are described in Supplementary Table 1. Participants were screened to ensure no history of neuropsychiatric disorders or systemic diseases. Participants were instructed to abstain from medications for 48 hours prior to assessment. Safety for TMS use was assessed using the Transcranial Magnetic Stimulation Adult Safety Screen (TASS) questionnaire (Keel et al., 2001). All original studies were conducted in accordance with the Declaration of Helsinki and were approved by the local ethics committees (N-20220018, N-20210047, N-20240004, N-20240053, N-2023040). Every participant provided written informed consent.

### 2.2. Transcranial Magnetic Stimulation Coupled with Electroencephalography

TMS-EEG data were acquired during single-pulse stimulation of the left M1. As this is a pooled dataset, acquisition parameters, including stimulator model and coil size, varied across the original studies (see Supplementary Table 1 for a complete breakdown by site). The M1 hotspot was defined as the scalp location eliciting the largest motor-evoked potential (MEP) or contraction from the contralateral first dorsal interosseous (FDI) muscle. Coil position was monitored using neuronavigation. TMS Evoked Potentials were assessed in real-time (Casarotto et al., 2022). Stimulation intensity was set at 90% of the resting motor threshold (rMT) and, in some subsets, it was adjusted to a minimum of 6*µ*V peak-to-peak. EEG data were recorded using a 64-channel, TMS-compatible g.HIamp amplifier (g.tec, Austria) with electrodes arranged according to the 10-20 system. Impedances were kept below 5 kΩ. Data were sampled at 4800 Hz and synchronized with TMS pulses.

### 2.3. Data Preprocessing

Data were preprocessed independently for each study using custom MATLAB or Python pipelines, as described in the original publications and summarized in Supplementary Table 1. Although specific algorithms (e.g., ICA, filtering) varied, all pipelines followed a comparable sequence: TMS artifact removal and interpolation; high-pass filtering and epoching around the TMS pulse; visual rejection of bad channels and noisy epochs; ICA to suppress ocular, muscular, and decay artifacts; spherical interpolation of removed channels; band-pass filtering; re-referencing to the common average; and baseline correction (−500 to −50 ms). All datasets were resampled to 300 Hz for group-level analysis. Additional details are provided in the Supplementary Table 1.

### 2.4. Feature Extraction

A comprehensive set of features (N=968) were extracted capturing response magnitude, oscillatory dynamics, and complexity. The core feature categories included:

i. Response Magnitude: Global/Local Mean Field Power (GMFP/LMFP - Skrandies, 1990) and Signal-to-Noise Ratio (GSNR/SNR - Luck, 2014).
ii. Spectral Dynamics: Global/Local Event-Related Spectral Perturbation (GERSP/ERSP - Cohen, 2014), Relative Spectral Power (GRSP/RSP - Cohen, 2014), and Natural Frequency (NF - Rosanova et al., 2009).
iii. Phase Consistency: Global/Local Inter-Trial Coherence (GITC/ITC - Cohen, 2014) and Phase-Locking Factor (GPLF/PLF - Aydore et al., 2013).
iv. Complexity: Perturbational Complexity Index using State Transition (PCIst - Comolatti et al., 2019).

Analyses were performed on all channels for global metrics and on predefined Regions of Interest (ROIs) for spatially specific (local) measures. Detailed mathematical definitions and algorithmic implementations references for each metric are provided in Supplementary Table 2-3.

### 2.5. Assessment of Group-Level Reproducibility using gTRCA

To corroborate that the large, heterogeneous normative group exhibited internally reproducible components, we applied group Task-Related Component Analysis (gTRCA) (Couto et al., 2026; Tanaka, 2020). Unlike standard averaging, gTRCA acts as a spatial filter to extract components that maximize within and between subject reproducibility. We validated the significance of these components against a null distribution of eigenvalues derived from 500 circle-shifted surrogates. Additionally, we assessed the stability of these components in the test-retest subset by correlating the spatial maps and time-series projections between sessions.

### 2.6. Reliability Assessment

Using the test-retest subset, we performed a comprehensive reliability analysis for all 968 features. To ensure a robust evaluation, we assessed three distinct aspects of reliability: systematic changes over time (i.e. systematic bias), the ability to distinguish between individuals (i.e. relative reliability), and the precision of the measurement scores (i.e. absolute reliability).

#### Systematic Bias

Data were evaluated for systematic bias to detect general drift or time effects between sessions. The normality of difference scores was assessed using the Shapiro-Wilk test. We applied paired-sample t-tests for normally distributed features and Wilcoxon signed-rank tests for non-normally distributed features. All p-values were corrected using the Benjamini-Hochberg False Discovery Rate (FDR) method. Features showing significant systematic bias (p < 0.05) were flagged, as this indicates a shift in the mean rather than random measurement error.

#### Relative Reliability

To assess the consistency of the *rank order* of subjects across sessions (i.e., how well the feature differentiates one participant from another), we calculated Relative Reliability. This was measured using the Intraclass Correlation Coefficient (*ICC*_{2,1}_; two-way random-effects model, absolute agreement; Koo & Li, 2016; Shrout & Fleiss, 1979), its 95% confidence interval (*ICC_↓_*_CI_, *ICC_↑_*_CI_) and Pearson’s correlation (*r*), or Spearman’s *p* for non-normal distributions. Reliability quality was assessed based on the lower bound of the 95% confidence interval of the *ICC*. Features were considered as having moderate (*ICC_↓_*_CI_ *T* 0.5), good (*ICC_↓_*_CI_ *T* 0.75), or excellent (*ICC_↓_*_CI_ *T* 0.9) reliability, provided they showed no significant systematic bias (*p*_FDR_ *T* 0.05).

#### Absolute Reliability

To quantify the magnitude of measurement error in the actual units of the feature (i.e., the precision of the score), we assessed Absolute Reliability. We calculated the Standard Error of Measurement (SM,; Cerin, 2023; Stratford & Goldsmith, 1997) and the Coefficient of Variation (CV%).

### 2.7. Data-Driven Feature Clustering

To visualize the structure of the high-dimensional feature space, we computed a correlation matrix and applied hierarchical clustering using Ward’s method (Ward, 1963). Based on the Scree plot, we identified three main clusters, using the merge distance to summarize the dataset’s variance and define the underlying dimensionality of the features.

### 2.8. Statistical Analyses

All analyses were performed using Python. We calculated descriptive statistics (mean, SD, median, 95% confidence interval (CI)) and assessed normality (Shapiro-Wilk test) for all features across the full cohort. As a substantial proportion of features deviated from a normal distribution, we reported median and interquartile ranges where appropriate and utilized non-parametric statistical tests for group comparisons when assumptions of normality were violated.

To isolate the normative physiological signal, we assessed age and sex effects using a two-step regression approach. First, an Interaction Model tested if age effects depended on sex (*Feature* ∽ *Age* x *Sex*). If no interaction was found, a Main Effects Model was fitted (*Feature* ∽ *Age* + *Sex*). Statistical significance was set at *p* < 0.05 after FDR correction.

### 2.9 Normative Comparison Tool (NormaTEP)

To enhance data accessibility while ensuring data privacy, we developed “NormaTEP,” an open-source, client-side web application. Using standard web technologies (HTML5, JavaScript) and statistical libraries (Math.js, jStat), the tool allows researchers to interactively filter the normative dataset by *Measure*, *Time*, *Cluster*, and *Band* to retrieve reference means (μ) and standard deviations (σ) and reliability outcomes. Beyond exploration, the tool includes a calculator for comparing single subject data against normative values. For univariate assessment, it computes standard Z-scores 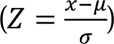 with visual indicators for values outside the 95% confidence interval. For multivariate assessment, the tool dynamically calculates the squared Mahalanobis distance (*D*^2^) for any user-selected combination of *k* features by subsetting the full covariance matrix (*Σ*) to construct a specific sub-matrix (*Σ*_sub_). The resulting distance, calculated as 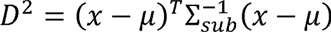, is evaluated against a Chi-squared distribution with *k* degrees of freedom to provide an immediate statistical assessment of global atypicality. To demonstrate the utility of this multivariate approach, we compared the normative values with new data from a healthy participant and a chronic pain patient. We defined a feature vector comprising three representative metrics, one selected from each identified cluster, based on previous reports on TMS-EEG and pain (De Martino et al., 2024, 2025; Jakobsen et al., 2025): alpha band (8–12 Hz) ITC and RSP, and broadband LMFP. All metrics were computed within the 15–120 ms time window in the left centro-parietal cluster, corresponding to left M1 stimulation.

## 3. Results

### 3.1 Dataset Description

A total of 164 healthy volunteers were included in the normative database. The cohort consisted of 88 females and 76 males, with a mean age of 30.8 ± 9.8 years (range: 20–71). The test-retest subset consisted of 28 females and 29 males, mean age: 28.5±6.1 (range: 21-48). Detailed demographic distributions and the specific contribution of each of the nine original studies are summarized in Table 1.

**Table 1.** Summary of demographic characteristics, TMS-EEG acquisition parameters, and source publications. This table details the composition of the normative dataset (N — 164; 88 female), including age (Mean *±* SD) and sex distribution for each contributing study. Methodological parameters for the primary motor cortex (M1) stimulation are provided, specifying the stimulator model, total pulse count, and Inter-Stimulus Interval (ISI). The “Publication” column refers to the original peer-reviewed studies from which the raw data were aggregated, with corresponding DOIs (when available) listed in the references below. Full description of the methodology per study can be found in the Supplementary Material. Studies marked with * were used in the test-retest subset (N=57; 28 female).

| *Study* | *N (Total/Female)* | *Age*<br>*(Mean ± SD)* | *TMS Stimulator* | *Number of*<br>*Pulses* | *ISI* | *Publication* |
| --- | --- | --- | --- | --- | --- | --- |
| 1 | 19 / 12 | 48.7 ± 13.3 | *MagPro R30* | 200 | 2 to 2.5 ms | (Jakobsen et al., 2025) |
| 2 | 22 / 12 | 28.7 ± 8.0 | *MagPro R30* | 200 | 2 to 2.5 ms | In prep. |
| 3 | 24 / 12 | 27.4 ± 5.5 | *MagStimRapid*<sup>2</sup> | 175 | 3 to 5 ms | (De Martino et al., 2024) |
| 4* | 12 / 4 | 31.8 ± 7.5 | *MagStimRapid*<sup>2</sup> | 200 | 2 to 2.5 ms | In prep. |
| 5* | 12 / 6 | 30.8 ± 6.8 | *MagStimRapid*<sup>2</sup> | 200 | 2 to 2.5 ms | In prep. |
| 6 | 18 / 9 | 29.4 ± 6.0 | *MagPro R30* | 200 | 2 to 2.5 ms | In prep. |
| 7 | 24 / 15 | 28.2 ± 4.8 | *MagStimRapid*<sup>2</sup> | 175 | 3 to 5 ms | In prep. |
| 8* | 20 / 12 | 24.6 ± 3.5 | *MagStimRapid*<sup>2</sup> | 175 | 3 to 5 ms | (Martino et al., 2024) |
| 9* | 13 / 6 | 29.3 ± 5.7 | *MagPro R30* | 200 | 2 to 2.5 ms | In prep. |
| **Total** | 164 / 88 | 30.8 ± 9.8 | - | - | - |  |

### 3.1 Spatiotemporal Characterization of Group-Level Responses

TEPs were extracted across the full cohort (*N* = 164). The GMFP heatmap revealed a highly consistent temporal structure across subjects, characterized by a rapid increase in activity at stimulus onset (*t* = 0 ms) followed by sustained power in the 0–250 ms window (**Figure 1A**).

**Figure 1.**
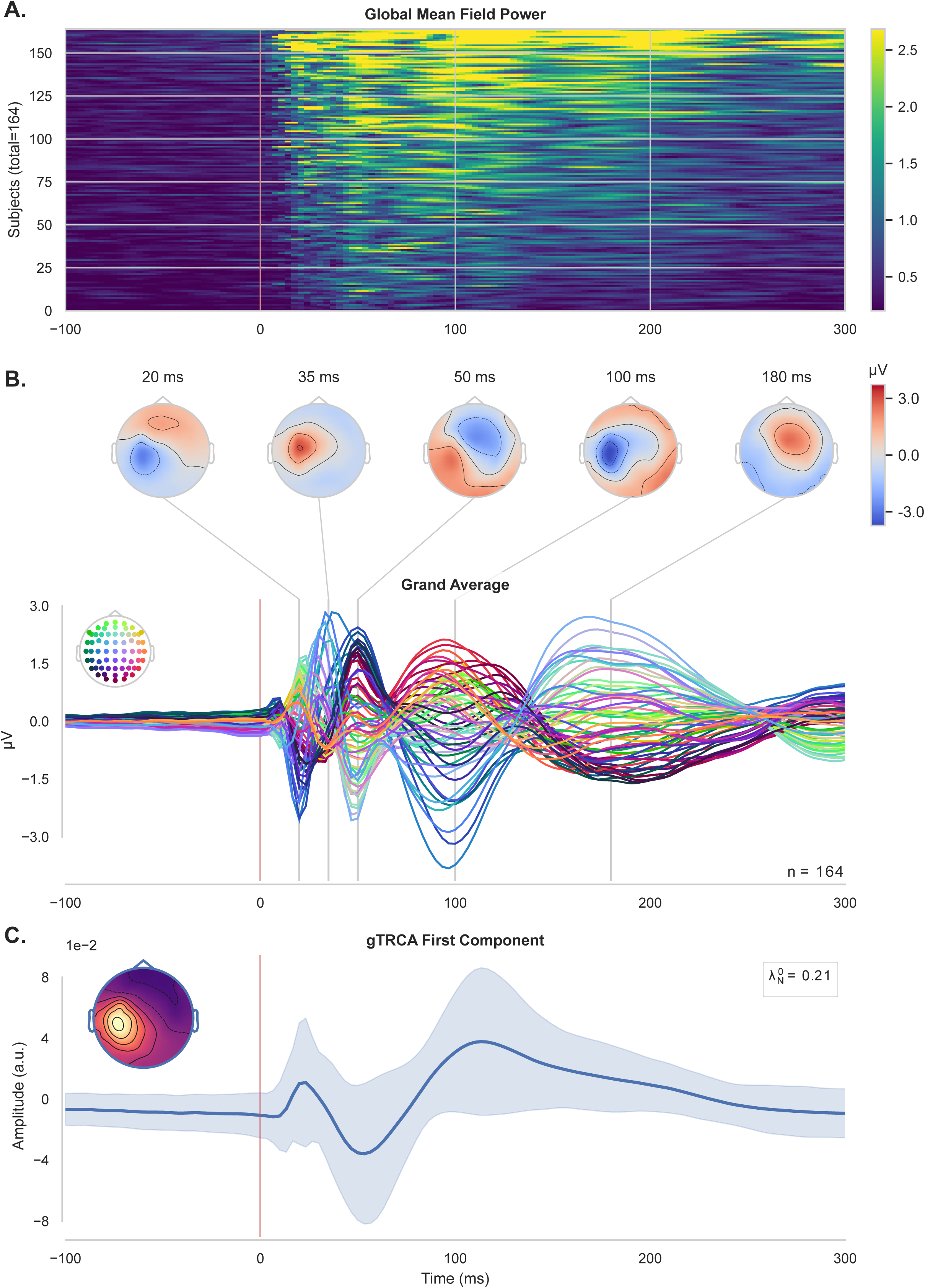
Group-level analysis of event-related responses and gTRCA component extraction (N — 164). (A) Heatmap of the Global Mean Field Power (GMFP) across individual subjects over time. Rows correspond to individual participants ordered by decreasing mean GMFP across the analyzed time (B) Grand average Event-Related Potentials (ERPs). The bottom panel shows a “butterfly plot” of grand averaged waveforms for all electrode channels (inset color legend indicates scalp location). The top row displays topographic voltage maps at selected peak latencies (20, 35, 50, 100, and 180 ms), illustrating the spatio-temporal evolution of the evoked response. (C) The first component extracted using group task-related component analysis (gTRCA). The main plot shows the component’s temporal waveform (mean ± 1.5 standard deviation indicated by shaded area), the normalized eigenvalue (2^0^), and the inset head map shows its corresponding spatial topography. This component significantly isolates the dominant underlying time-locked task-related neural pattern shared within and between subjects in the normative dataset (p<0.002).

The grand average TEPs, visualized via a butterfly plot of all electrode channels, confirmed a robust evoked response (**Figure 1B**) showing similar characteristics to previous reports (Beck et al., 2024; Couto et al., 2026; Fecchio et al., 2025; Ni et al., 2022; Tremblay et al., 2019). Distinct voltage topographies emerged at specific latencies, suggesting a sequence of activation shifting from posterior to anterior regions. Key polarity reversals were observed at approximately 20, 35, 50, 100, and 180 ms. Importantly, gTRCA yielded two components significantly reproducible at the group level (**Supplementary Figure 1**). The temporal profile of the dominant first component is characterized by a sharp biphasic deflection between 20–50 ms and a focal left centro-parietal topography consistent with the TMS coil positioning (**Figure 1C**).

### 3.2 Test-Retest Reliability

TEPs spatiotemporal structure remained highly consistent across test-retest sessions, with GMFP heatmaps (**Figure 2A**) and TEP butterfly plots (**Figure 2B**) mirroring the profiles observed in the normative cohort (cf. **Figure 1A, B**). The gTRCA eigenspectra were similarly stable, consistently identifying two significant components (**Supplementary Figure 1, right panel**). The first component’s strength was virtually unchanged between sessions 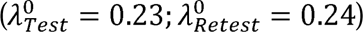 (**Figure 2C**). Visual inspection of the extracted component time series and spatial filters confirmed high qualitative agreement (**Figure 2C; cf. Figure 1C**). This was quantitatively supported (**Figure 2D**) by strong subject-wise correlations (*r* = 0.71 ± 0.23) and stable spatial topographies (*r* = 0.66 ± 0.29).

**Figure 2.**
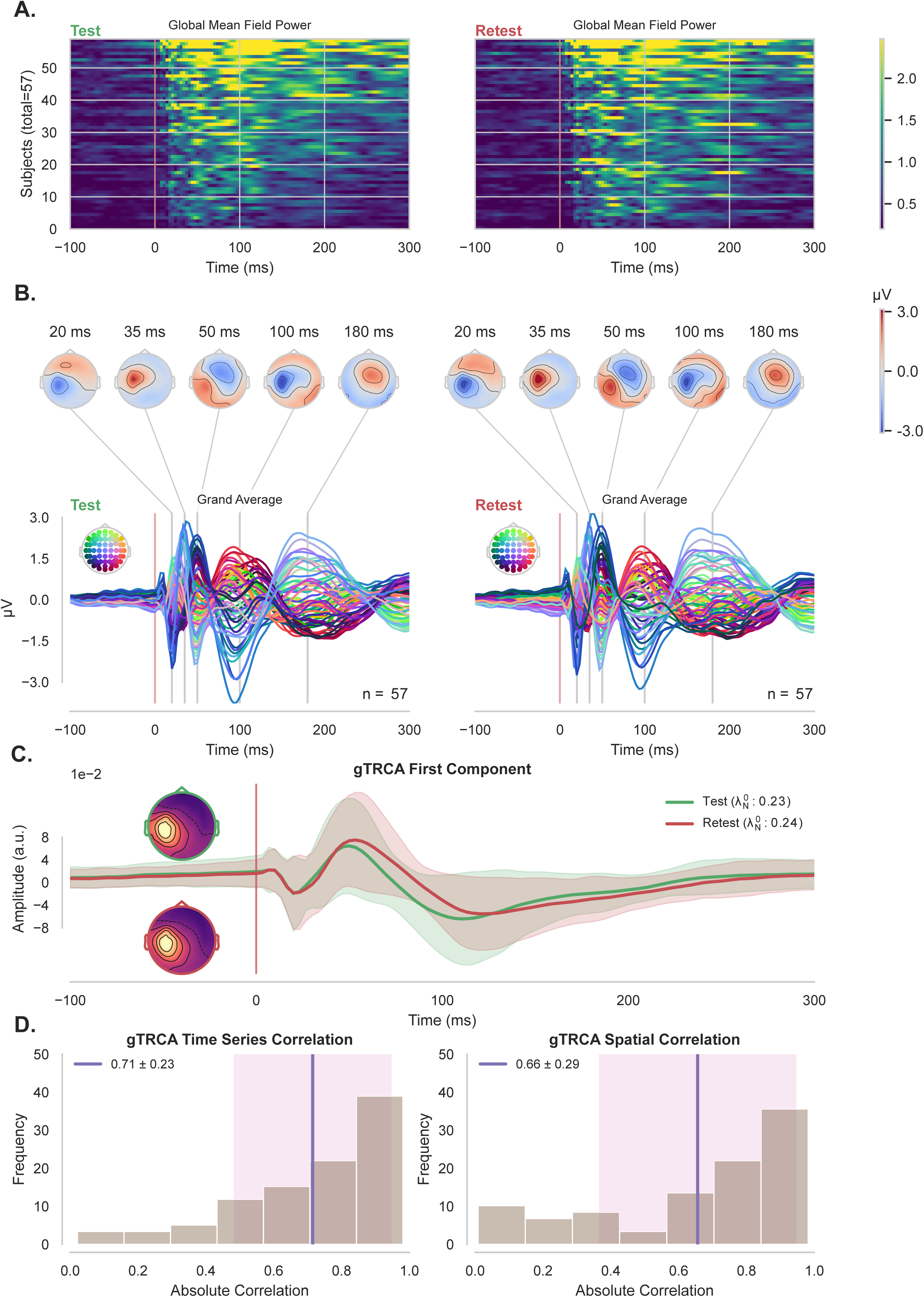
Test-retest reliability of event-related responses and gTRCA components (N — 57). (A) Global Mean Field Power (GMFP) for individual subjects during Test (left) and Retest (right) sessions. Heatmaps show consistent response magnitudes across the cohort in both sessions. (B) Comparison of Grand Average ERPs. Butterfly plots display the superimposed electrode waveforms for Test and Retest conditions. Topographic maps at 20, 35, 50, 100, and 180 ms illustrate the reproducibility of spatial voltage distributions over time. (C) The first component extracted via gTRCA for Test (green) and Retest (red) sessions. The temporal waveforms (mean ± 1.5 standard deviations) and inset spatial topographies demonstrate high overlap and consistency between sessions. (D) Quantitative reliability assessment. Histograms display the frequency distribution of absolute subject-wise correlations for the gTRCA time series (left; *r* — 0.71 *±* 0.23) and spatial topographies (right; *r* — 0.66 ::: 0.29) between Test and Retest sessions (purple lines indicate the mean and shaded region indicates the standard deviation).

### 3.3 Evaluation of Feature Reliability

No systematic bias was identified for the vast majority of features (99.27%), with 95% confidence interval of the mean difference including zero. Absolute reliability was also high across the feature space, with a median CV of 18.1% (95% CI: 15.2% – 21.0%).

As shown in **Figure 3**, relative reliability varied across analytical dimensions and was generally moderate (ICC 0.5) across time windows, with higher values in the early (15–120 ms) compared to the late (180–300 ms) window (**Figure 3A**).

**Figure 3.**
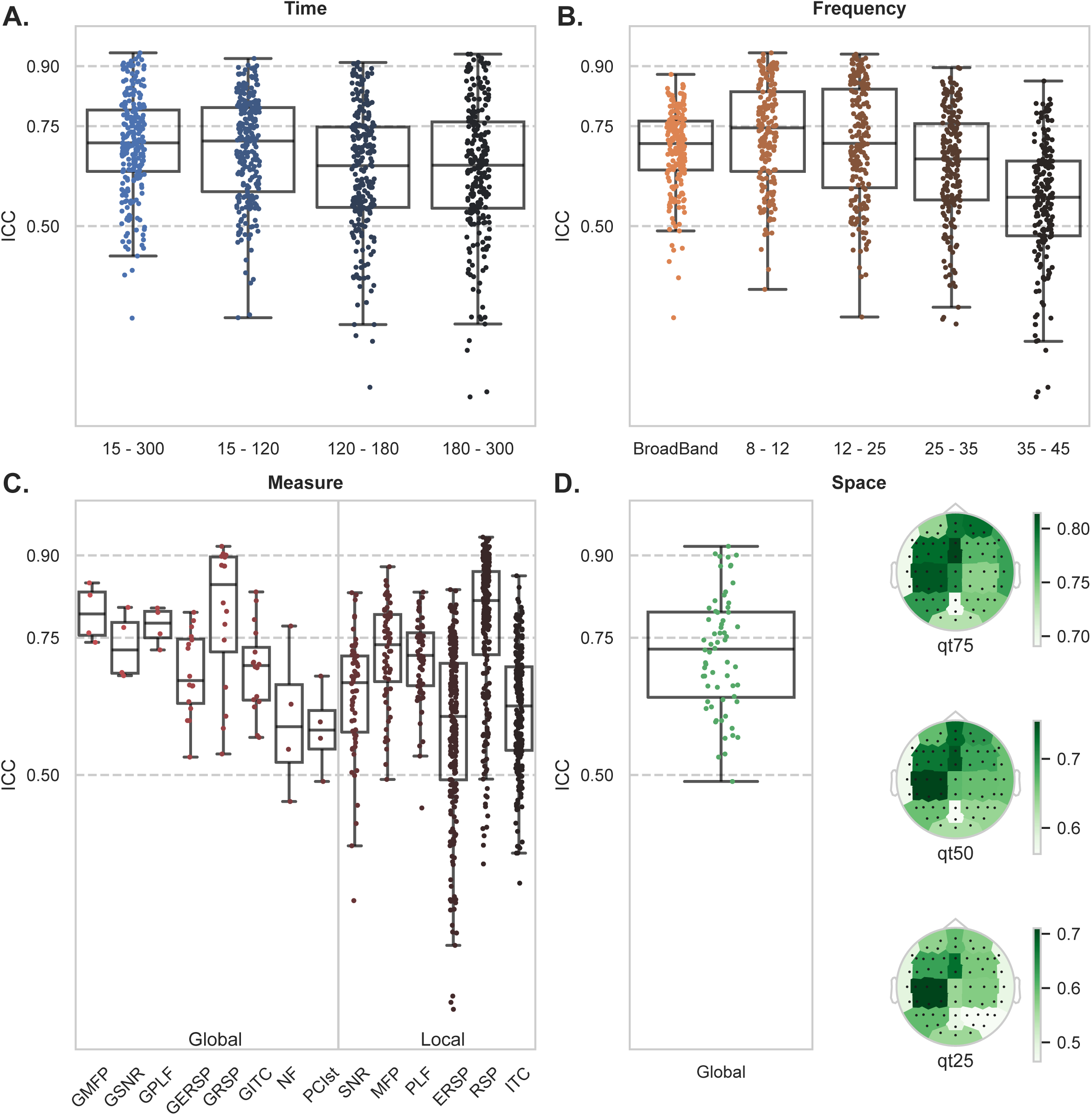
Intraclass Correlation Coefficient (ICC) values across different feature parameters. Box plots display the distribution of ICC values (y-axis) with individual data points overlaid, evaluating test-retest reliability across four dimensions: (A) Time Windows: Reliability assessed across four temporal intervals (15–300 ms, 15–120 ms, 120–180 ms, and 180–300 ms). (B) Frequency: Reliability across Broadband and specific frequency ranges (8–12 Hz, 12–25 Hz, 25–35 Hz, and 35–45 Hz). (C) Measures: Comparison of reliability for Global (left) and Local (right) metrics. Global measures include GMFP, GSNR, GPLF, GERSP, GRSP, GITC, NF (only computed under the stimulation site), and PCIst while Local measures include SNR, MFP, PLF, ERSP, RSP, and ITC. (D) Space: A summary of global reliability (left) and the spatial distribution of ICC values across the scalp (right), illustrated by topographic maps at different quantiles (qt25, qt50, qt75).

Broad-band features and lower frequency bands (8–25 Hz) demonstrated highest median reliability. A noticeable decline in ICC values was observed in the Gamma band (35–45 Hz), consistent with the lower signal-to-noise ratio typically observed in high-frequency EEG components (**Figure 3B**). Most global metrics outperformed many local metrics in terms of reliability (**Figure 3C**). Specifically, ERSP and ITC showed a wide range of reliability depending on the specific set of channels and the frequency probed. Topographic mapping of ICC values across all local features revealed a clear spatial gradient, with the highest reliability clusters (*ICC* > 0.75) located in the left central and parietal regions, while peripheral sensors showed lower stability (**Figure 3D**). The full reliability profile for all 968 features, including ICC, SEM and bias results, can be found in the NormaTEP web application https://boutoo.github.io/NormaTEP/.

### 3.4 Unsupervised Clustering of Feature Architecture

Hierarchical clustering of correlations between TMS-EEG features (**Figure 4A**) identified a three-cluster solution, as indicated by a sharp “elbow” in the scree plot, with the merge distance dropping from 100% to 22.1% (**Figure 4B**).

**Figure 4.**
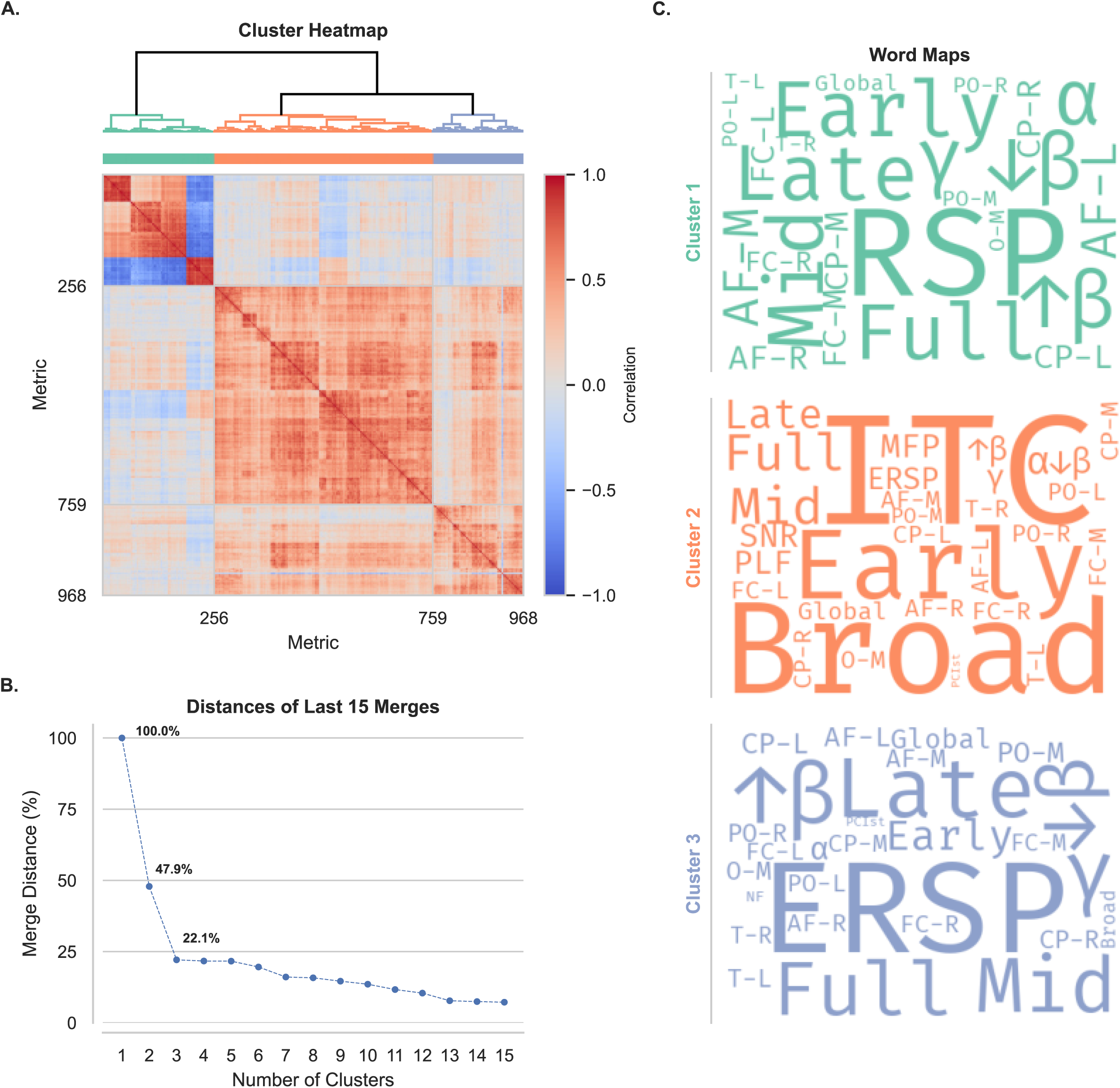
Hierarchical clustering and characterization of feature correlations. (A) Cluster Heatmap. The matrix displays pairwise correlations between all extracted features. Hierarchical clustering was performed using Ward’s method, resulting in a dendrogram (top) that organizes the features into three distinct groups (color-coded green, orange, and blue). (B) Cluster optimization. The scree plot shows the merge distances for the final 15 merges. The distinct “elbow” at 3 clusters (22.1% distance) validates the decision to partition the data into three main groups. (C) Feature Word Maps. Word clouds visualize the composition of each cluster (Cluster 1: Green; Cluster 2: Orange; Cluster 3: Blue). The size of each word represents the frequency with which that term appears in the feature labels assigned to that cluster, highlighting the dominant metrics, time windows, bands and clusters characterizing each group. A list with all label’s definitions can be found in the Supplementary Material.

Word frequency maps (**Figure 4C**) showed that the three clusters were differentiated primarily by the type of signal metrics computed, rather than by temporal divisions. Qualitatively, cluster 1 (Green) was dominated by the term RSP alongside a mix of temporal labels (“Early,” “Mid,” “Late”), identifying it as a group representing normalized spectral power distribution independent of specific time windows. Cluster 2 (Orange) was defined by terms such as ITC and “Broad” (Broadband), reflecting phase-consistency and synchronization across frequencies. Cluster 3 (Blue) was mainly characterized by ERSP, capturing absolute power modulations and magnitude changes and other distinct metrics. Average intra-cluster correlation values where 0.644 ± 0.195, 0.482 ± 0.150, and 0.390 ± 0.193 for Cluster 1, 2, and 3 respectively.

### 3.5 Normative Modeling and Single-Subject Outlier Detection

Finally, we integrated all features into a multivariate normative model. Shapiro-Wilk tests indicated that most raw features did not follow a Gaussian distribution (*p* < 0.05), validating the need for non-parametric or transformed modeling approaches (**Figure 5A**).

**Figure 5.**
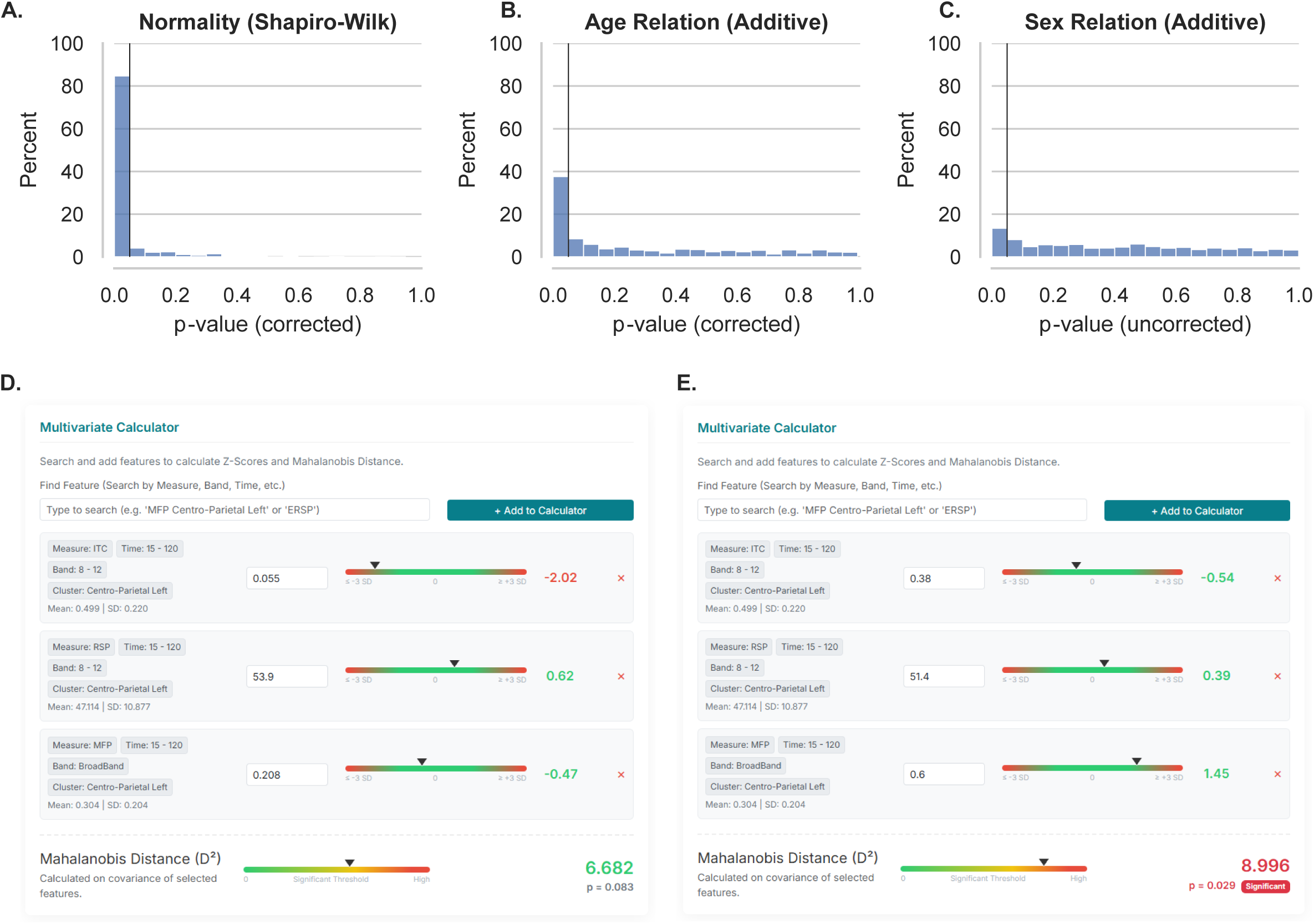
Normative modeling and multivariate outlier detection. (A) Assessment of univariate normality using the Shapiro-Wilk test. The distribution of corrected p-values indicates that most features deviate from a normal distribution (skew towards 0). (B-C) Evaluation of demographic confounders. Histograms show the distribution of p-values and vertical line indicates p — 0.05 for (B) Age (FDR-corrected) and (C) Sex (uncorrected) relationships using an additive model. (D-E) Multivariate abnormality assessment tool using NormaTEP web application for (D) new healthy participant and (E) a chronic pain patient. The red-green-red bars display the z-score (≤—3 to ≥ 3 standard deviations) for each assessed feature (namely the early centro-parietal left ITC, RSP and MFP) with each z-value on the right side. The bottom green-yellow-red bars displays the Mahalanobis distance (D^2^ score) with its associated value on the right.

The impact of demographic covariates on feature variance was further assessed. An additive model revealed a significant relationship between Age and feature values (skewed corrected p-values, **Figure 5B**), whereas Sex had a negligible effect (non-skewed p-value distribution, **Figure 5C**).

To illustrate clinical applicability, we projected two test cases into this normative space. The first healthy profile (**Figure 5D**) exhibited a univariate outlier (z-score < 2) in a single feature but was correctly classified as normative by the integrated index, albeit approaching the threshold for significance (*D*^2^ = 6.682, *p* = 0.083). Conversely, the chronic pain patient (**Figure 5E**) showed no univariate deviations (all |*z*| < 2) yet was objectively flagged as abnormal in the multivariate analysis (*D*^2^ = 8.996, *p* = 0.029).

## 4. Discussion

This study establishes a comprehensive, reliability-tested normative database of TMS-EEG metrics, representing a critical step toward the clinical translation of TMS-EP-based excitability and connectivity assessment. By analyzing a large-scale cohort with a wide array of metrics, we provide an initial map of cortical excitability and connectivity in symptom-free human adults.

### Reliability and Stability

The core contribution of this work lies in the systematic quantification of test-retest reliability across nearly 1,000 TMS-EEG features. We found that 54.3% of extracted metrics met rigorous moderate-to-high relative reliability standards. Notably, we adopted a deliberately conservative methodological stance: rather than relying on ICC point estimates, we categorized features based on the lower bound of the 95% confidence interval. This choice ensures that any metric labelled as ‘good’ or ‘excellent’ reflects a highly probable minimum level of reproducibility, accounting for the inherent estimation uncertainty in smaller samples. In a field where even a second-decimal fluctuation can shift an index, this restrictive criterion is very conservative and prevents unintended overestimation of stability.

Our results build upon and extend recent reliability assessments (e.g., Bertazzoli et al., 2025). While previous work highlighted the challenges of obtaining reliable early-latency components due to lower signal-to-noise ratio in smaller datasets, our findings demonstrate that these components can be highly stable (ICC > 0.59) when assessing larger sample sizes. This suggests that the early evoked response (15–120 ms) is a robust physiological marker when artifacts are managed through the large-scale harmonization and rigorous cleaning pipelines employed here (Ziemann et al., 2026). Conversely, the ∼45% of features that failed to meet these criteria were not random; they were predominantly late-window spectral measures (e.g., Gamma and high-Beta ITC and ERSP). This suggests that while early evoked components are highly stable, late-oscillatory responses may reflect more transient physiological states or be more susceptible to signal-to-noise limitations and might require more trials or specific processing to be used as biomarkers.

Beyond relative consistency, we found negligible systematic bias across the database, with most features showing no significant drift between sessions, and exhibiting good-to-excellent CV values. Notably, the median upper bound of the CV (*CV*_95_) remained near the 20% threshold, suggesting that the majority of TEP features provide sufficient measurement precision for individual-level longitudinal tracking.

### Feature Space and Dimensionality Reduction

The clustering analysis revealed a structured organization of the TMS-EEG feature space, which we propose not as a definitive taxonomy, but as a functional guide for dimensionality reduction. The identification of non-overlapping signal clusters is essential for moving toward leaner, more focused clinical protocols that avoid the “fishing expeditions” common in high-dimensional neurophysiological data.

### From Univariate Outliers to Multivariate Profiles

A pivotal shift proposed here is the move from group-level univariate Z-scores to individualized multivariate profiles. Traditional approaches that tag “abnormal” metric features one-by-one inherently suffers from a high false-positive rate when dealing with large number of metrics. A multivariate lens treats an individual’s response as a single integrated signature. This approach was illustrated by our clinical case example, which demonstrated a double dissociation: we identified a chronic pain patient as abnormal despite having no single outlier feature, and a healthy control as “normal” despite one univariate outlier. Furthermore, our analysis identified age as a primary driver of variance across multiple features, whereas sex showed negligible influence, a finding described for motor evoked potential-based connectivity measures (Cueva et al., 2016), that adds clarity to the ongoing debate regarding demographic impacts on TEPs, while also highlighting age-correction as a needed step for normative modelling, as done here.

### Stability Across Platforms and Limitations

Remarkably, the neural components identified were stable across data acquired with different stimulator brands and coil sizes. While the data were aggregated from existing studies (convenience sampling), a limitation that could introduce methodological bias, the fact that harmonized cleaning pipelines yielded such reproducible results across hardware validates the robustness of the “perturb-and-measure” logic. Another important observation is that our current database is skewed toward younger adults (20–71 years, mean age 30.8), which may limit precision in geriatric populations. Additionally, the focus on the left M1 means these “fingerprints” may not generalize to other targets like the DLPFC. Following the ENIGMA consortium model (Quidé et al., 2022), the database is designed to be expandable, with future iterations aiming to include more diverse demographics and non-motor cortical regions.

## 5. Conclusion

This work provides a critical basis for a community-wide effort. The resulting comprehensive normative database, supported by the developed open-access web interface, provides an immediate tool for researchers to compare their findings and assess M1 TMS-EEG features expected reliability. By integrating reliability mapping with multivariate normative modeling, this study lays the foundation for moving TMS-EEG from a descriptive experimental technique to a quantitative clinical tool.

## Acknowledgements

We are grateful to Teis Kalsbøl Nielsen, Kristian Eskildsen and Vithushan Amirthanathan for their assistance with TMS-EEG data collection. We also thank Dennis Boye Larsen for his insightful discussions regarding the visual representation of the data and figure design. Finally, the authors thank Kenia Fuentes for her careful review of the manuscript and feedback on its clarity and accessibility.

## Author Contributions

BANC: Conceptualization, Methodology, Software, Formal Analysis, Visualization, Writing – Original Draft.

E.D.M: Investigation, Writing - Review & Editing.

D.M.S: Methodology, Software, Formal Analysis, Writing - Review & Editing.

A.J.: Investigation, Writing - Review & Editing.

M.M.B: Investigation, Writing - Review & Editing.

A.G.: Investigation, Writing - Review & Editing.

S.I.M.: Investigation, Writing – Review & Editing.

T.G.N.: Conceptualization, Methodology, Writing - Review & Editing, Supervision.

A.G.C.: Conceptualization, Methodology, Software, Validation, Formal analysis, Investigation, Resources, Data Curation, Writing - Original Draft, Writing - Review & Editing, Visualization, Supervision.

D.C.A.: Conceptualization, Methodology, Formal analysis, Data Curation, Writing - Review & Editing, Visualization, Supervision, Project administration, Funding acquisition.

## Data Availability Statement

The normative distributions and feature reliability scores generated during this study are integrated into the NormaTEP web application, which is publicly accessible at https://boutoo.github.io/NormaTEP/. The underlying raw data from the constituent studies are available from the corresponding author of the related publications upon reasonable request, subject to the data-sharing agreements of the contributing institutions.

## Declaration of Generative AI and AI-assisted Technologies

During the preparation of this work the author(s) used AI for language editing.

## SUPPLEMENTARY MATERIAL

**Supplementary Figure 1.** Identification of significant gTRCA components using surrogate data testing. Plots display the eigenspectrum (λ_N_) of components extracted via group Task-Related Component Analysis (gTRCA) against a logarithmic x-axis. Significance thresholds (dashed horizontal lines) were established using the maximum eigenvalues from 500 circle-shifted surrogate datasets. (Left) Results for the full normative dataset (N — 164). Two components exceeded the noise floor defined by the surrogates (λ_N_ > 0.147 — 2), indicating they contain significant task-related signals. (Right) Results for the test-retest subset (N — 57). The eigenvalue distribution is highly reproducible between the Test (green circles) and Retest (red squares) sessions. Both sessions consistently yield two significant components above the chance level (λ_Ntest_ > 0.166 — 2 and λ_Nretest_ > 0.165 — 2).

**Supplementary Table 1.** Specific data acquisition parameters per dataset. Extra details are described below. Studies marked with asterisk were also used in the test-retest subset. All studies used a figure-of-eight coil.

**Supplementary Table 2.** Mathematical definitions and computational implementations of the EEG feature space.

**Supplementary Table 3. Summary of TMS-EEG feature extraction parameters and permutations.** Measures are categorized by computational domain and spatial scale (Global vs. ROI-specific). The table defines the integration of four time windows and four frequency bands across 15 predefined spatial clusters to derive the multidimensional feature space.

