## Supplementary figures and images for "The Rhythm of Normality: A Comprehensive Normative Database for TMS-EEG Metrics with Reliability Characterization"

### Supplementary Figure 1

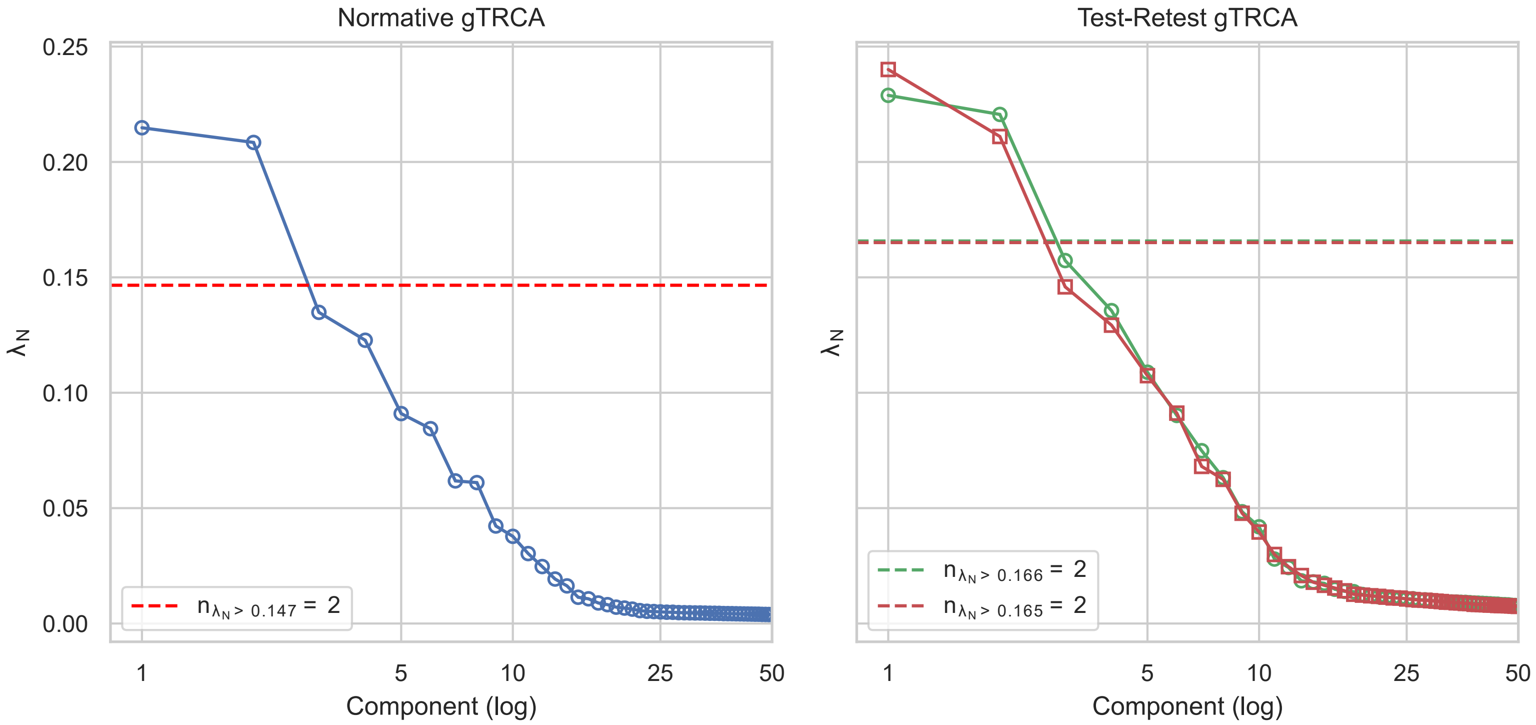
