## Supplementary Table 1 for "The Rhythm of Normality: A Comprehensive Normative Database for TMS-EEG Metrics with Reliability Characterization"

| Study | Original Study | Stimulator & Coil | Coil Size | Waveform | Intensity | Auditory Masking | EEG System | Preproc. Software | Filter Settings |
| --- | --- | --- | --- | --- | --- | --- | --- | --- | --- |
| 1 | (Jakobsen et al., 2025) | $MagPro R30$ | 65 | Biphasic | 90% rMT | Foam earplugs + TAAC | g.HIamp (64ch) | Python (MNE) | 1-80 Hz |
| 2 | *In prep.* | $MagPro R30$ | 65 | Biphasic | 90% rMT | Foam earplugs + TAAC | g.HIamp (64ch) | Python (MNE) | 1-80 Hz |
| 3 | (De Martino et al., 2024) | $MagStimRapid^{2}$ | 70 | Biphasic | 90% rMT (~$6\mu V PtP$) | Foam earplugs + TAAC | g.HIamp (64ch) | MATLAB (EEGLAB) | 0.5-80 Hz |
| 4* | *In prep.* | $MagStimRapid^{2}$ | 70 | Monophasic | 90% rMT (~$6\mu V PtP$) | Foam earplugs + TAAC | g.HIamp (64ch) | Python (MNE) | 0.5-80 Hz |
| 5* | *In prep.* | $MagStimRapid^{2}$ | 70 | Biphasic | 90% rMT (~$6\mu V PtP$) | Foam earplugs + TAAC | g.HIamp (64ch) | Python (MNE) | 1-80 Hz |
| 6 | *In prep.* | $MagPro R30$ | 65 | Biphasic | 90% rMT | Foam earplugs + TAAC | g.HIamp (64ch) | Python (MNE) | 1-80 Hz |
| 7 | *In prep.* | $MagStimRapid^{2}$ | 70 | Biphasic | 90% rMT | Foam earplugs + TAAC | g.HIamp (64ch) | Python (MNE) | 1-80 Hz |
| 8* | (Martino et al., 2024) | $MagStimRapid^{2}$ | 70 | Biphasic | 90% rMT (~$6\mu V PtP$) | Foam earplugs + TAAC | g.HIamp (64ch) | MATLAB (EEGLAB) | 0.5-80 Hz |
| 9* | *In prep.* | $MagPro R30$ | 65 | Biphasic | 90% rMT | Foam earplugs + TAAC | g.HIamp (64ch) | Python (MNE) | 1-80 Hz |
| Description | | | | | | | | | |
| Across nine studies, navigated TMS-EEG was performed using a 64-channel g.HIamp system. While the majority of studies utilized a biphasic pulse, Study 4 was unique in its use of a monophasic waveform. Hardware configurations were split between two primary setups: a M200² (Magstim Co., UK) stimulator with a 70mm coil (Studies 2, 3, 4, 5, 6, 7, 9) and an MagVenture R30 (MagVenture A/S, Denmark) stimulator with a 65mm coil (Studies 1 and 8). Studies 2, 3, 4, 6, 7, 9 used Brainsight (Rogue Research Inc., Montreal, QC, Canada) and studies 1, 5, 8 used Invesalius (Renato Archer Information Technology Center, Campinas, SP, Brazil) for real-time coil tracking. The reference electrode was placed on the right mastoid (A2) for all studies, except study 9, which placed it between the eyebrows.  Stimulation Intensity was set at 90% rMT for all datasets. The resting motor threshold (rMT) was determined via MEP in four studies (2, 3, 7, 9) and through visual inspection in the remaining studies. Stimulation intensity was further adjusted to a minimum of $\boldsymbol{6\mu V}$ peak-to-peak on channels under the coil in five studies (2,3,6,7,8). Pulse counts were largely standardized at 200, though Studies 3 and 7 utilized 170 pulses, and Study 9 utilized 175 pulses.  Offline preprocessing followed two software-specific pipelines, using MNE Python (studies 1, 4, 5, 8, and 9) or Matlab custom scripts (studies 2, 3, 6, and 7). Studies 1, 4, 5, 8, and 9 utilized MNE-Python. Both pipelines followed a similar structure: TMS artifact removal (-2 to 15 ms), visual trial and bad channels inspection/rejection, ICA-based cleaning of muscle, ocular, and decay artifacts. Bad channels were interpolated and the signal was set to average reference. Finally, band-pass filtering was applied at 1–80 Hz, with Studies 3, 4 and 8 extending to 0.5–80 Hz. | | | | | | | | | |

**Supplementary Table 1.** Specific data acquisition parameters per dataset. Extra details are described below. Studies marked with asterisk were also used in the test-retest subset. All studies used a figure-of-eight coil.
