## Supplementary Table 2 for "The Rhythm of Normality: A Comprehensive Normative Database for TMS-EEG Metrics with Reliability Characterization"

| Measure | Implementation Logic | Formal Definition |
| --- | --- | --- |
| Mean Field Power | Root Mean Square (RMS) of voltage across all $K$ channels. | $\boldsymbol{GMFP}\left( t \right)=\sqrt{\frac{1}{K}\sum_{i=1}^{K} V_{i}\left( t \right)^{2}}$ |
| Signal-to-Noise Ratio | Ratio of Root Mean Square amplitude between response and baseline windows. | $\boldsymbol{SNR}=\sqrt{\overline{V_{resp}^{2}}/\overline{V_{base}^{2}}}$ |
| Phase Locking Factor | Hilbert transform applied to broadband signal; instantaneous phase ($\phi$) consistency across trials (n). | $\boldsymbol{PLF}\left( t \right)=\left\vert\frac{1}{N}\sum_{n=1}^{N} \phi_{c,n}\left( t \right) \right\vert$ |
| Inter Trial Coherence | Morlet wavelet convolution derived phase (w) consistency across trials (n) at specific frequencies. | $\boldsymbol{ITC}(f, t)=\frac{1}{N}\sum_{n=1}^{N} \frac{W_{n}\left( f,t \right)}{\left\vert W_{n}\left( f,t \right) \right\vert}$ |
| Event-Related Spectral Perturbation | Baseline-corrected spectral power using a log-ratio (dB) transformation. | $\boldsymbol{ERSP}\left( f,t \right)=10\log_{10} \left( \frac{P\left( f,t \right)}{\mu_{base}\left( f \right)} \right)$ |
| Relative Spectral Power | Summed power in band divided by total spectral power. | $\boldsymbol{RS}\boldsymbol{P}_{band}=\frac{\sum P_{band}}{\sum P_{total}}\times100$ |
| Natural Frequency | Frequency of peak aggregated power in the early TFR window. | $\boldsymbol{NF}=\text{argmax}_{f}\left( \sum_{t} P_{f,t} \right)$ |
| Perturbational Complexity Index - State Transition | State transitions in the principal components of the TEP. | $\boldsymbol{PC}\boldsymbol{I}^{\boldsymbol{ST}}=\sum_{n} \Delta NST_{n}$ |
| Observations | | |
| To reduce the multidimensional data into discrete features for analysis, different aggregation strategies were applied based on the metric's domain. Time-domain magnitude and phase consistency metrics (MFP, PLF) were quantified as the Area Under the Curve (AUC) using the trapezoidal rule across specified time windows ($\boldsymbol{t}_{\boldsymbol{lim}}$), capturing the total integrated response.  Conversely, spectral dynamics (ERSP, ITC) were calculated by averaging values across the respective frequency bands ($\boldsymbol{f}_{\boldsymbol{lim}}$) and time windows ($\boldsymbol{t}_{\boldsymbol{lim}}$). Relative Spectral Power (RSP) was expressed as a percentage ratio of band-specific power to total broadband power (8–45 Hz). Complexity (PCIst) and Natural Frequency (NF) were extracted as discrete scalar values representing state transition density and peak oscillatory frequency, respectively.  Unlike Inter-Trial Coherence (ITC), which was calculated in the frequency domain, PLF was derived by applying a Hilbert transform to the broadband signal to extract the analytic phase. To ensure the robustness of the PLF, a Rayleigh-based statistical comparison was implemented. A null distribution was estimated from the pre-stimulus baseline, and a Rayleigh scale parameter was derived. Only phase-locking values exceeding the $\boldsymbol{1-}\boldsymbol{\alpha}_{\boldsymbol{corr}}$ threshold (where $\boldsymbol{\alpha}$ was FDR-corrected for the number of channels) were included in the final AUC calculation.  Metrics are reported using a prefix notation where 'G' indicates a global average across the entire scalp, and its absence indicates a local calculation within a given spatial ROIs. | | |

**Supplementary Table 2.** Mathematical definitions and computational implementations of the EEG feature space.
