## Supplementary Table 3 for "The Rhythm of Normality: A Comprehensive Normative Database for TMS-EEG Metrics with Reliability Characterization"

| **Measure** | **Domain** | **ROIs (Clusters)** | **Time Windows (ms)** | **Frequency Bands (Hz)** |
| --- | --- | --- | --- | --- |
| **Global Measures** | | | | |
| GMFP | Time | Global | All 4 | N/A |
| GSNR | Time | Global | All 4 | N/A |
| GPLF | Phase | Global | All 4 | N/A |
| GERSP | Time-Frequency | Global | All 4 | All 4 |
| GRSP | Time-Frequency | Global | All 4 | All 4 |
| GITC | Phase (Time-Frequency) | Global | All 4 | All 4 |
| NF | Time-Frequency | Only CP-L | All 4 | N/A |
| PCIst | Complexity | Global | All 4 | N/A |
| **ROI-Specific Measures** | | | | |
| SNR | Time | All 15 Spatial ROIs | All 4 | N/A |
| MFP | Time | All 15 Spatial ROIs | All 4 | N/A |
| PLF | Phase | All 15 Spatial ROIs | All 4 | All 4 |
| ERSP | Time-Frequency | All 15 Spatial ROIs | All 4 | All 4 |
| RSP | Time-Frequency | All 15 Spatial ROIs | All 4 | All 4 |
| ITC | Phase (Time-Frequency) | All 15 Spatial ROIs | All 4 | All 4 |
| **Parameter** | **Total Count** | **Sub-categories** | | |
| Time Windows | 4 | • Early: 15 - 120 ms  • Mid: 120 - 180 ms  • Late: 180 - 300 ms  • Full-Time (Full): 15 - 300 ms | | |
| Frequency Bands | 4 | • Alpha ($\alpha$): 8 - 12 Hz  • Low Beta ($\downarrow\beta$): 12 - 25 Hz  • High Beta ($\uparrow\beta$): 25 - 35 Hz  • Gamma ($\gamma$): 35 - 45 Hz  • Broad-band (Broad): 8 – 12 Hz | | |
| Regions of Interest (ROIs) | 15 | • Anterior-Frontal (AF): Left, Midline, Right (3)  • Fronto-Central (FC): Left, Midline, Right (3)  • Temporal (T): Left, Right (2)  • Centro-Parietal (CP): Left, Midline, Right (3)  • Parieto-Occipital (PO): Left, Midline, Right (3)  • Occipital (O): Midline (1) | | |

**Supplementary Table 3. Summary of TMS-EEG feature extraction parameters and permutations.** Measures are categorized by computational domain and spatial scale (Global vs. ROI-specific). The table defines the integration of four time windows and four frequency bands across 15 predefined spatial clusters to derive the multidimensional feature space.
